# Automated Morphometric Analysis Reveals Plasticity Induced by Chronic Antidepressant Treatment in Hippocampal Astrocytes

**DOI:** 10.1101/2020.11.26.399964

**Authors:** Parul Sethi, Garima Virmani, Surya Chandra Rao Thumu, Narendrakumar Ramanan, Swananda Marathe

## Abstract

Nervous system development and plasticity involves changes in cellular morphology, making morphological analysis a valuable exercise in the study of nervous system development, function and disease. Morphological analysis is a time-consuming exercise requiring meticulous manual tracing of cellular contours and extensions. We have developed a software tool, called SMorph, to rapidly analyse the morphology of cells of the nervous system. SMorph performs completely automated Sholl analysis. It extracts 23 morphometric features based on cell images and Sholl analysis parameters, followed by Principal Component Analysis. SMorph is tested on neurons, astrocytes and microglia and reveals subtle changes in cell morphology. Using SMorph, we found that chronic 21-day treatment with antidepressant desipramine results in a significant structural remodeling in hippocampal astrocytes. Given the proposed involvement of astroglial structural changes and atrophy in major depression in humans, our results reveal a novel kind of structural plasticity induced by chronic antidepressant administration.

## Introduction

Cells of the nervous system are characterized by intricate morphology. Precisely built cytoarchitecture allows precise cell-to-cell contacts between different cells, which is indispensable for their function ^1^. For example, neurons are highly compartmentalized cells with complex dendritic trees and precisely guided axon terminals, which allow the neurons to form synapses with their specific pre- and postsynaptic partners ^2^. Stimuli that profoundly affect the brain function and behavior, such as chronic stress, epileptic seizures, stroke and neurological disorders, lead to marked changes in neuronal structure ^3-7^. Furthermore, the process of neurogenesis during development and adulthood involves rapid changes in dendritic arborization ^8,9^ which necessitates careful quantitative analysis of dendritic arbors.

Additionally, other cell types in the Central Nervous System (CNS), including astrocytes, also possess complex cellular morphology characterized by precise contacts with synaptic compartments ^10^. They have been shown to be highly plastic in response to various physiological and pathological stimuli, thus aiding synaptic plasticity and function ^11^. In particular, astrocytes play a key role in synaptic transmission and plasticity ^12-14^. Perisynaptic astrocytic processes (PAPs) show activity-dependent motility ^15^. Induction of long term potentiation (LTP) stabilizes the PAPs, which in turn results in spine stability ^15^. Astrocytes are also instrumental in the pathology related to trauma, inflammation and degeneration ^16,17^. A significant feature of astrocytes in these pathologies is a process called reactivation, which involves copious morphological remodeling along with molecular, cytoskeletal and functional changes, which ultimately play a key role in the disease outcome ^10,18-21^. On the other hand, under physiological conditions, astrocytes may undergo subtler changes in morphology, which may potentially bring about significant changes in synaptic physiology and behavior ^11^,^15^,^22^. Especially, changes in astrocytes in major depression have received particular interest. Studies in depressed patients consistently show reduced astrocyte density in key limbic brain regions ^23–25^. Additionally several studies in animal models of depression have shown reduced glial density and morphological complexity ^26-30^, which may be reversed by chronic antidepressant treatment ^26^. Interestingly, depletion of prefrontal astrocytes has been shown to be sufficient to induce depressive symptoms in experimental rats ^31^, thus suggesting a causal link between glial loss and behavioral symptoms of major depressive disorder (MDD). In spite of the growing evidence, the morphological alterations in astrocytes in response to stress and antidepressant treatments have not been studied in detail.

Owing to the complex and diverse shapes that astrocytes assume *in vivo*, it has been challenging to analyze and quantify the subtle morphological changes in astrocytes. Moreover, manual analysis of one cell at a time greatly limits the sample sizes and number of experimental groups that can be analyzed at once. Sholl analysis is a valuable tool to quantify the morphological complexity of nervous system cells, where the number of processes emanating from the cell body are counted as a function of the distance from the centroid ^32^. While most Sholl analysis protocols rely on manual neurite tracing, tools do exist that allow completely automated Sholl analysis on single cells ^33,34^. However, tools to perform Sholl analysis on large datasets with no manual intervention are particularly scarce. Another major drawback of Sholl analysis is that it may not reveal subtle changes in dendritic arborization. Hence, tools are needed for a more comprehensive and sensitive analysis of cellular morphometry in a multiparametric space. Here, we present a method that works on neurons, astrocytes and microglia. This method, called SMorph, allows for unbiased, completely automated investigation of cellular morphology. As a proof of principle, we analyzed the morphology of astrocytes following a unilateral stab wound injury and compared it to the astrocytes on the contralateral side. Performance of SMorph was compared to the existing methods that allow some degree of automation. SMorph performed 1080 times faster than the fastest method tested, without compromising the accuracy. In addition, it also performs the complete multiparametric morphometry unlike any other method followed by PCA.

Using SMorph, we asked if chronic treatment with an antidepressant, desipramine, in mice results in morphological changes in astrocytes. Our analysis showed a yet unknown form of structural plasticity in astrocytes as a result of chronic antidepressant treatment. Such plasticity may help reverse the adverse changes in astrocytic morphology observed in depressed patients and in animal models of depression. We believe that SMorph would help uncover structural changes happening in the cells of the nervous system in an expeditious and unbiased manner on large datasets.

## Materials and Methods

### Animal experiments

All animal experiments were carried out in accordance with the guidelines from the IISc institutional animal ethics committee (IAEC). Experiments were carried out on C57BL6/J mice in IISc central animal facility. Mice were housed in groups of 3-5 mice per cage, unless stated otherwise, and were maintained on a 12-12 hour light-dark cycle with access to water and food *ad libitum*. 4-5 month old adult male mice (25-32g) were used in the study. All efforts were made to reduce the number of animals by using the principle of 3R’s.

### Stab wound surgery

Stab wound experiment was performed as described previously with minor modifications ^35^. In brief, mice were anesthetized with 3% isoflurane (Sorane, NEON) and mounted on a stereotax (KOPF, USA) with continuous administration of 1.5-2% isoflurane with air until the end of the surgical procedure. Head was shaved and the skin was disinfected with the betadine solution. Midline incision was made and the skin was removed. A hole was made on the skull on the right side of the sagittal suture at 1.8 mm caudal and 2 mm lateral of bregma. A 2 mm deep stab injury was made using a #11 scalpel blade mounted on the stereotax (Fig. S1). The scalpel was held in place for 30 seconds and retracted gently. Afterwards, the skull was covered using acrylic dental cement. After completion of the surgical procedure, mice were transferred onto a heat pad maintained at 40°C for 5-10 minutes until the mice woke up. 3 mice were used for the stab wound experiment. Following the surgery, mice were single-housed. 4 days after the stab wound procedure, mice were sacrificed by transcardial perfusion with saline solution (0.9% NaCl) followed by 4% paraformaldehyde (PFA) and the brain was dissected out. The tissue was processed as described in the following section.

### Desipramine treatment

For chronic antidepressant treatment, 5 mice received 20 mg/kg desipramine intraperitoneally (i.p.) daily for 21 consecutive days (Fig. 3 A). 5 control animals received equivalent volume (10 ml/kg) of 0.9% saline solution for the same period. 2 hours after the last drug injection, mice were sacrificed by transcardial perfusion with saline solution followed by 4% paraformaldehyde (PFA). Brains were dissected out and post-fixed overnight at 4°C in 4% PFA. Brains were cryo-protected by placing them in 30% sucrose solution in PBS until they sank to the bottom. Thereafter, the brains were frozen in blocks of OCT solution (Sigma) and kept at −80°C until sectioning. Serial floating coronal sections (40μm thick) were cut using a cryostat (Leica, CM 1850) and were stored at 4°C in 0.1M Phosphate Buffer (PB) until staining.

**Figure 3:**
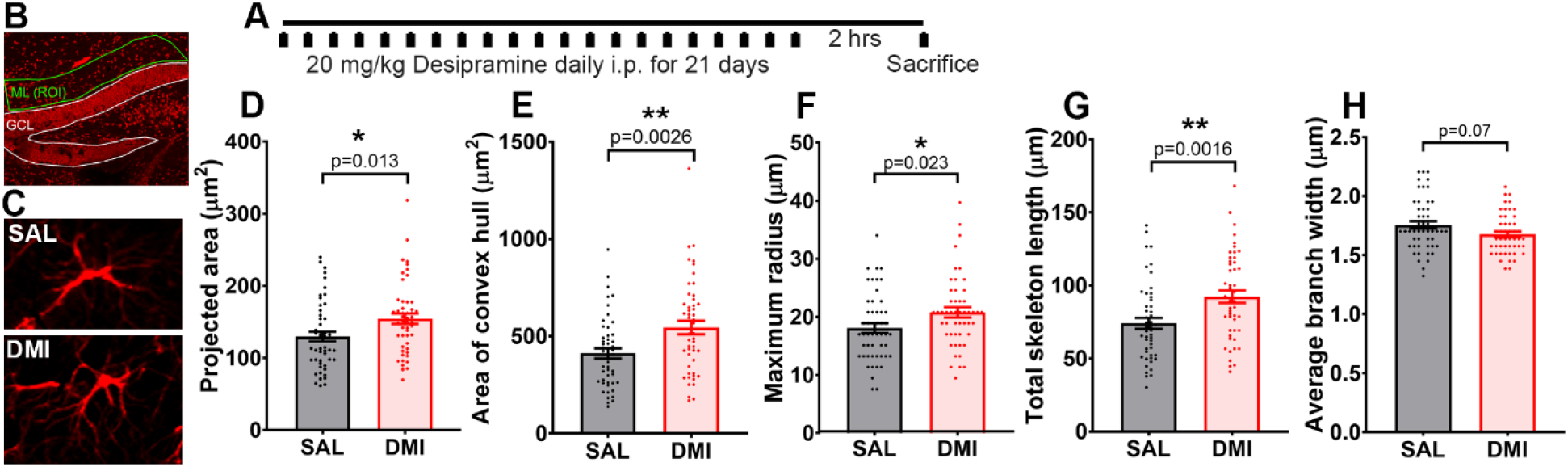
Chronic antidepressant treatment produces subtle increase in parameters related to cell size. **(A)** The schematic represents the experimental paradigm depicting that the mice were treated intraperitoneally with 20mg/kg of desipramine (DMI) once daily for 21 consecutive days. Control mice received an equivalent volume of 0.9% saline solution. Mice were sacrificed by transcardial PFA perfusion 2 hours after the last drug injection. **(B)** Representative epifluorescence image (DAPI in red) shows the ROI, comprising the molecular layer (ML) above the upper blade of the dentate gyrus granule cell layer (GCL), from where GFAP-immuno positive astrocytes were cropped. Representative examples of astrocytes cropped from **(C)** controls and **(D)** DMI-treated mice are shown. Bar graphs with scatter points show a statistically significant increase in the **(D)** 2D-projected area, **(E)** area of the convex Hull, **(F)** maximum radius and **(G)** total skeleton length, in DMI-treated mice as compared to controls. **(H)** Bar graph with scatter points shows a trend towards decline in the average branch width in the DMI-treated mice as compared to the controls. Bars represent mean±SEM. Asterisks in D-G represent statistically significant differences, with corresponding p values shown. n= 50 cells per group, evenly sampled from 5 mice per experimental group.

### Immunohistochemistry and imaging

Immunohistochemistry was performed as described before ^27^. In brief, sections were washed 3 times with PB and blocked with 10% normal donkey serum + 3% Bovine Serum Albumin (BSA) + 0.3% TritonX100 in 0.1M PB at room temperature for 1 hour. Following this, sections were incubated with a primary antibody overnight at 4°C (chicken pAb anti-GFAP; 1:1000; Novus, NBP1-05198). Following the incubation with the primary antibody, sections were washed 3 times with PB and then incubated with the secondary antibody for 2 hours (goat anti chicken Alexa Fluor 594, abcam, ab150172). After incubation with the secondary antibody, sections were washed with PB 3 times. Sections were mounted on glass slides in the mounting medium with DAPI (Abcam ab 104139). The slides were imaged on a Zeiss LSM 880 airyscan confocal microscope with a Plan-Apochromat 20X/0.8 M27 objective. 1024X1024 pixels, 8-bit images were acquired at 1 μm z-step size. For stab wound experiments, the ROI was selected in the penumbra region, slightly away from the site of the wound. A similarly placed ROI from the contralateral side was used as a control (Fig. S1). For the saline vs desipramine comparison, the molecular layer of the upper blade of the dentate gyrus (DG) was used as an ROI (Fig. 3 B). Image metadata was used to convert distances in pixels to μms manually for representation in the desipramine experiment.

### Image analysis and software development

For morphological analysis, single cells were cropped out in X-Y and Z dimensions and 2-D maximum intensity projection (MIP) images were obtained (Fig. 1 A-C). These steps were performed manually using ImageJ. However, these steps can also be automated using machine learning tools published elsewhere ^36,37^. Images of cells from 2 different experimental groups were saved into 2 separate folders named appropriately. Further steps were completely automated. The output data is generated as.txt files and it auto-populates the folders named “Sholl results”, “Features” and “pca values”. The high-intensity blobs were detected as a position of soma, to be used later for Sholl analysis and the calculation of maximum radius (Fig. 1 D). Boundary blobs coming from adjacent cells were discarded from analysis. Following this, the images were thresholded to obtain a binary image containing only the high-intensity pixels, and the background was removed by selecting the largest continuous trace, while all other traces were removed (Fig. 1 D). Sum of all pixels was taken at this stage to obtain the total 2D-projected surface area of a cell. Afterwards, the thresholded image was converted to a skeletonized image by successively removing boundary pixels, unless it broke the continuity of the object, until process width was reduced to 1 pixel (Fig. 1 E). Sum of all pixels was taken at this stage to obtain total skeleton length. The ratio of the 2-D projected area to the total skeleton length was calculated to obtain average branch width. We then moved the detected soma position to its nearest point on the skeleton. Following this, the astrocyte branches from the skeletonized image were classified into classes such as primary, secondary, tertiary, quaternary, and terminal, based on their position respective to the estimated location of the centre of the soma or the extremities and the branch points (Fig. 1 F). Primary, secondary, tertiary, quaternary branches were labeled with red, blue, pink and green colors respectively. Other higher order branches, if present, were classified together into one class and labelled cyan (Fig. 1 F). Convex hull was drawn around the extremities of the cell skeleton and its area was estimated in pixels (Fig. 1 G). For Sholl analysis, concentric circles were drawn around the soma position with successive radii increasing by 3 pixels (Fig. 1 H). We plotted the number of intersections of the skeletonized cell as a function of the radius (Fig. 1 I). We used polynomial regression to obtain final plots for Sholl analysis for individual cells. Polynomial-fitted Sholl plots for all the cells in a given experimental group were expressed as mean±SEM.

**Figure 1:**
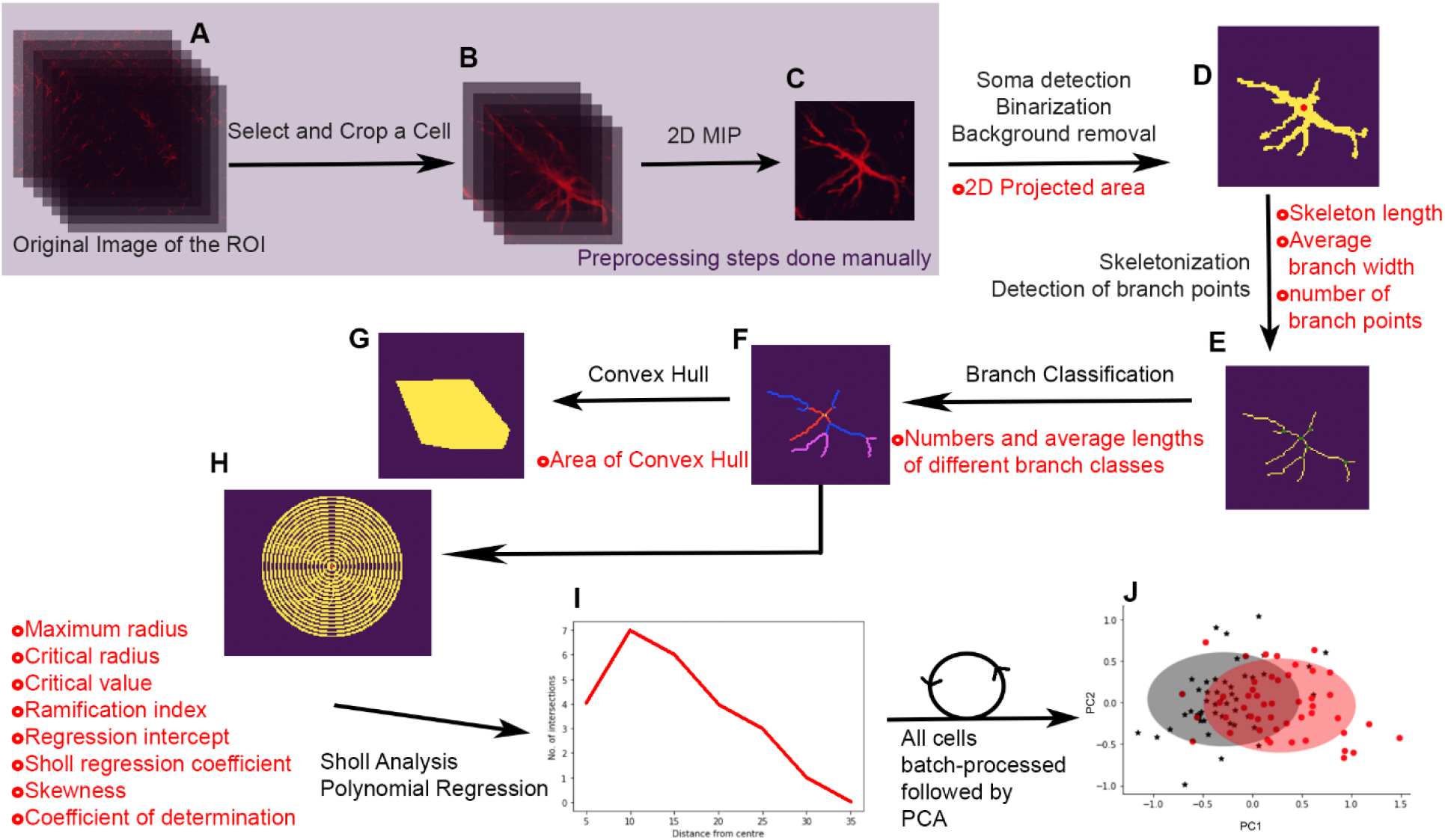
Image processing and analysis pipeline. **(A)** Confocal images of the ROI are used to select the cells to be analyzed. **(B)** The selected cells are cropped in X, Y and Z dimensions. **(C)** z-stacks of individual cells are flattened into 2 dimensional maximum intensity projections. These 3 steps are performed manually on ImageJ. **(D)** Soma was detected using blob detection. Peripheral blobs belonging to adjacent cells were discarded. Images of single cells were then thresholded to obtain binary images. Following this largest contiguous trace was selected and the background was removed. **(E)** The binary silhouettes of cells were then skeletonized. The detected soma was moved to the nearest point on the skeleton. The branching points were detected based on the sum of nearest neighboring pixels. **(F)** Branches were classified based on their position with respect to the estimated soma position and/or extremities. Here, primary, secondary and tertiary branches are labeled red, blue and pink respectively. **(G)** Convex hull was drawn around the extremities of the cell skeleton to calculate the area of the convex hull. **(H)** concentric circles with successive radii increasing by 3 pixels were drawn around the soma and **(I)** number of intersections were plotted as a function of the distance from the soma. Features extracted at each step are listed in red. **(J)** All these steps are repeated on each cell to generate a feature vector for every cell, which is then used to perform PCA.

Following feature were extracted from the images of single cells: total length of skeleton, average branch width, 2D-projected area, area of convex hull, maximum enclosing radius, numbers and average lengths of primary, secondary, tertiary, quaternary and terminal branches, number of branch points, critical value (local maximum of the polynomial fit), critical radius (radius corresponding to the critical value), schoenen ramification index (ratio between critical value and number of primary branches), regression intercept, Sholl regression coefficient (rate of decay of no. of branches), Skewness, and coefficient of determination (of the linear regression of the data). All these parameters were then collectively used as feature vectors for PCA (Fig. 1 J).

The cross-platform image processing pipeline was written in python 3 using the “scikit-image” library and all of the above steps were automated. The pre-processing steps for blob detection, thresholding and skeletonization were implemented using scikit-image library’s Laplacian of Gaussian, Otsu threshold and skeletonize functions ^38^. Cubic polynomial fitting for Sholl plots of individual cells was implemented using scikit-learn’s polynomial feature generation ^39^ followed by linear regression ^38^. PCA implementation provided in the decomposition module of scikit-learn was then used to infer the first 2 principal components based on features extracted from our data ^39^. The feature vectors were scaled using MaxAbsScaler from the scikit-learn’s preprocessing module before being used for PCA so as to prevent any sub-optimal results. First two principal components were plotted as 2 dimensional scatter plots and were overlaid with ellipses representing 3 standard deviations around the means. The complete code is available in the supplemental zip file (SMorph.zip).

### Installation

For analysis with SMorph, we recommend the use of UNIX-based operating systems such as Linux or MacOS. Nevertheless, if windows users face any issues, please follow the troubleshooting instructions from https://github.com/parulsethi/Morphometric-analysis. The code has been tested in python 3.7, which can be downloaded from https://www.python.org/downloads/release/python-370/. Install the dependencies by typing the following in the command line: “pip install-r requirements.txt”. The repository is available in the supplemental zip file (SMorph.zip). To perform analysis on your own datasets, place your data folders inside the Morphometric analysis folder and implement analysis by replacing the input data folder names. To run the notebooks, execute “jupyter notebook” from the command line and locate the desired notebook using your browser. The notebook single_cell_analysis.ipynb includes visual analysis to explore various morphological parameters of a single cell. Notebook group_analysis.ipynb includes the group level analysis of astrocytes using Principal Component Analysis (PCA) which helps to distinguish the differences between different classes of astrocytes based on their morphological parameters. The cropped cells need to have a good contrast ratio for automated batch processing. Representative individual cells can be tested with the single cell analysis notebook (single_cell_analysis.ipynb) to determine suitability of images for the fully automated analysis. Small adjustments to contrast ratios may be required in some cases. If the batch processing stops midway, it is likely due to the image quality of a specific cell. Please check the last image name on the list shown in the group analysis Jupyter notebook and either enhance the contrast ratio of that cell or delete that cell, and rerun the program.

### Manual image analysis

We compared the performance of SMorph with the analysis performed using the built-in functions and external plugins in ImageJ. Images were thresholded to obtain binary images followed by background removal using the “analyse particle” tool. Remaining background, if any, was deleted manually. Pixel sum was taken at this stage to obtain the projected 2D-surface area of the cells. Following this, the images were skeletonized using the “skeletonize” plugin in ImageJ. Pixel sum taken at this stage gave the total skeleton length of the cells. The ratio of the area to skeleton length was calculated to obtain the average pixel width. To calculate the area of the convex hull, all the extremities of the cell skeletons were marked manually using the point selection tool in ImageJ. The built-in convex hull function was used to convert point selections into an area that represented the smallest convex hull around the cells. Afterwards, the “measure” function was used to calculate the area of the selection. Number of branch points and the number of terminal branches were obtained using the “analyse skeleton” tool in ImageJ. Linear regression through the origin was performed between values obtained through ImageJ and SMorph. R-squared (R^2^) was used to determine the correlation.

To perform Sholl analysis, we used 2 different methods involving ImageJ to optimize for either speed or accuracy. In one method, skeletonized images obtained as described above were used. However, we observed that the skeletons thus obtained in an unsupervised manner contained several artefacts, the most prominent being the closed loop-like structures along the processes, which rendered the analysis error-prone. To perform a more accurate Sholl analysis, manual process tracing was performed by a trained blind observer using ImageJ to obtain skeletonized cells. This was followed by Sholl analysis using the ImageJ plugin ^34^. The centre of the soma was selected as the centre for Sholl analysis. ImageJ macro scripts were used wherever possible to perform batch-processing in order to minimize the time taken. In all cases, the total time taken for the analysis was monitored using a stopwatch, and rounded off to the closest multiple of 5 minutes. This was compared to the total runtime for SMorph. The comparison between SMorph and ImageJ was performed on 46 cells of the control group in the stab wound experiment.

### Statistical analysis

All statistical analyses were performed using GraphPad Prism 8.0.2 and the results were expressed as mean ± SEM. For comparisons of morphology-related parameters between groups, statistical outliers were determined using the ROUT method at Q=1%, and were removed from analysis. The normality of the datasets was determined using Kolmogorov-Smirnov normality test with Dallal-Wilkinson-Lilliefor approximation. The distribution was deemed to be normal (Gaussian) at the alpha of 0.05. Two group comparisons were made using t-test to determine statistical significance in case of normal distribution or by Mann-Whitney U test whenever the data was not normally distributed. For Sholl analysis, multiple comparisons were made with multiple t-tests, using the false discovery rate (FDR) approach, with a 2-stage step-up method of Benjamini, Krieger and Yekutieli. The differences between groups were deemed to be significant after the discovery was determined at Q = 1%.

## Results

### Stab injury results in significant structural changes in CA1 stratum oriens astrocytes

In order to assess if SMorph can be used to detect and quantify the structural remodeling in astrocytes, we performed a unilateral stab injury to the *stratum oriens* region of hippocampal CA1 (Fig. S1). Stab wound is known to induce robust astroglial reactivation resulting in substantial structural remodeling ^40^. We analyzed the morphology of astrocytes on the ipsilateral side and compared it to those on the contralateral side as a control. A total of 46 cells from the control side and 48 cells from the stab wound side evenly sampled from 3 mice were used for the analysis (Fig. 2 A-D). We found a significant increase in the parameters related to cell size; viz., total skeleton length, average thickness of the processes, 2D-projected surface area occupied by the cells, area of the convex hull drawn around the extremities of the cells, and the maximum enclosing radius (Fig. 2 E). We also observed a statistically significant increase in the parameters related to branching of the astrocytic processes on the stab wound side, as compared to the contralateral side. These parameters include number of tertiary branches, average length of tertiary branches, number of branch points and number of terminal branches (Fig. 2 E). Interestingly, we did not find any change in the numbers and average lengths of the primary and secondary branches (Fig. 2 E), which suggests that stab wound-induced astroglial reactivation mainly induces branching at the distal ends, while branches close to the soma are largely unaffected. We further performed Sholl analysis which revealed a significant increase in the ramification of astrocytes on the ipsilateral side as compared to the contralateral side (Fig. 2 F). We also found a statistically significant increase in the area under the Sholl curve (Fig. 2 G). A total of 23 features related to the astrocytes morphology were extracted from images and the feature vector was used for PCA (Fig. 2 H). The first two principal components described about 50% of the observed variability in the data sets. The first principal component correlated positively with the size and branching complexity of the astrocytes. We found a statistically significant increase in the PC1 inferred for the astrocytes from the ipsilateral side as compared to that for the contralateral side (Fig. 2 I). This analysis showed that SMorph can be used to rapidly compare the morphology of astrocytes between experimental groups. Further, we inferred that the profuse branching occurred more in distal branches as compared to the proximal branches. This data suggested that such detailed analysis of cell morphology can also be used to rapidly identify subtle changes in astrocyte morphology induced by physiologically relevant stimuli in a completely automated and unbiased manner.

**Figure 2:**
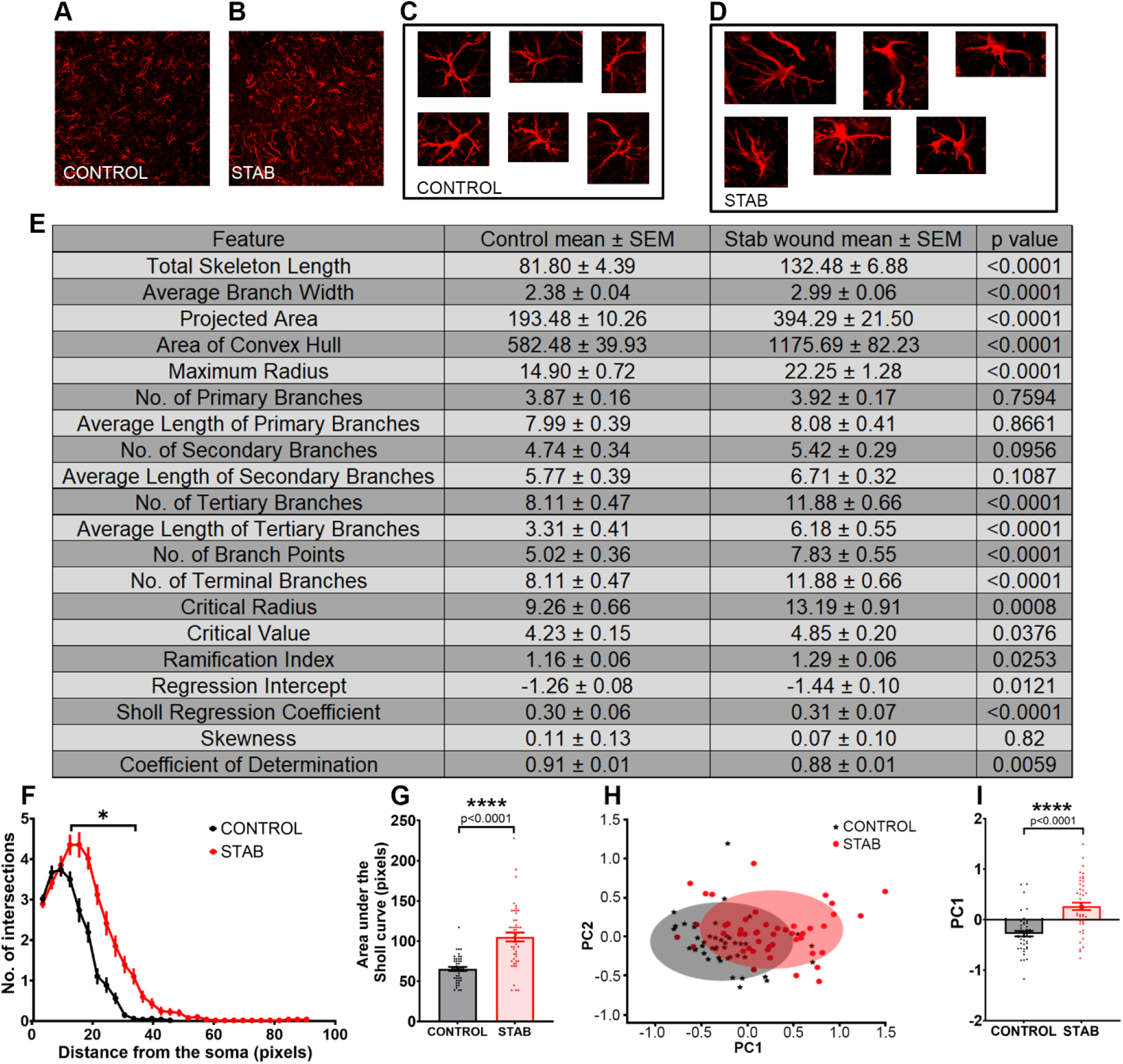
Stab injury results in significant structural changes in CA1 stratum oriens astrocytes. Mice underwent a unilateral stab wound surgery targeting the CA1 *stratum oriens*. ROI was selected from the penumbra region near the stab wound. Matching ROI from the contralateral side was used as a control. Shown are the representative images of ROIs from **(A)** the controls and **(B)** the stab wound images. Representative images of cropped GFAP-immunopositive astrocytes from **(C)** controls and **(D)** stab wound are shown. **(E)** Table shows mean±SEM of individual features extracted from images of astrocytes from controls and stab wound. The table also shows p-values comparing the 2 groups. **(F)** The line plot shows a significant increase in ramification in the stab wound group as compared to the controls in Sholl analysis. **(G)** Bar graph overlaid with scatter points of area under the Sholl analysis curve shows a significant increase in the stab wound group as compared to the controls. **(H)** 2-dimensional scatter plot depicts the first two principal components with ellipses corresponding to 3 standard deviations around the means. **(I)** Bar graph with scatter points depicts a significant increase in PC1 in the stab-treated mice as compared to the controls. Bars represent mean±SEM. Asterisk in F represents discovery using the FDR approach. Asterisks in G and I represent statistically significant differences, with corresponding p values shown. n= 46 cells (Controls) and 48 cells (Stab wound) evenly sampled from 3 mice.

### SMorph allows quick and accurate morphometry analysis

We tested the performance of SMorph against the analysis performed using built-in functions and plugins in ImageJ. Custom ImageJ macro scripts were used to automate several successive image processing steps on the entire batch of images, wherever possible. Time taken for these analyses was monitored using a stopwatch and was compared to the total runtime for SMorph on the same batch of cells. We computed the 2D-projected area, total skeleton length and average process width of the cells using built-in functions in ImageJ. We found a strong correlation between the values calculated by SMorph and those by ImageJ. For the total 2D-projected area of cells, we found an R^2^ value of 0.97, and the slope of 0.83 (Fig. S2 A). The R^2^ for the total skeleton length was estimated to be 0.97, and corresponding slope of the linear regression was 0.84 (Fig. S2 B). In the case of average branch width, the R^2^ value was found to be 0.99, whereas the slope of the line was 0.98 (Fig. S2 C). It took 2 hours and 35 minutes to calculate the area, 55 minutes to calculate the skeleton length and 10 minutes to compute the average branch width. We then measured the area of the convex hull manually using built-in ImageJ functions and found an R^2^ value of 0.95 and slope of 0.83 between SMorph and ImageJ estimates (Fig. S2 D). It took 3 hours and 5 minutes in total to perform the calculation of the area of the convex hull. We further calculated the number of branch points and the number of terminal branches using the “analyse skeleton” plugin in ImageJ. We found an R^2^ value of 0.89 with slope of 0.60 for the number of branch points (Fig. S2 E), while the corresponding values for the number of terminal branches were 0.92 and 0.99 respectively (Fig. S2 F). The analysis of the number of branch points in ImageJ led to a consistent overestimation because of the closed-loop structures that were generated as artefacts of unsupervised skeletonization, which accounted for a lower value of slope (Fig. S2 E). It took a total of 1 hour and 55 minutes to perform these two analyses.

To perform Sholl analysis as quickly as possible, we used an ImageJ macro to batch-process the process of skeletonization followed by Sholl Analysis plugin ^34^. This method slightly overestimated the number of processes, mainly because of the closed-loop artifacts in skeletons generated in an unsupervised manner. Hence, we also performed completely manual tracing of astrocytic processes followed by Sholl analysis ^34^ for a more accurate measurement of the number of processes. We determined that the Sholl analysis with neither SMorph nor semi-automated ImageJ was significantly different from the results of the completely manual analysis (Fig. S2 G). While the analysis through completely manual tracing took 7 hours and 40 minutes in total, batch processing using ImageJ macro could be completed within 3 hours and 20 minutes.

SMorph computed more parameters than our ImageJ analysis and performed PCA, within only 40 seconds (Fig. S2 H). The output of SMorph was comparable to the manual analysis using ImageJ. Furthermore, SMorph also provides additional parameters that none of the other methods provides, which makes it possible to further analyze the morphometric features of cells in a multidimensional space.

We reanalysed the control vs stab images for sholl analysis using the semi-automated ImageJ method and found that the stab wound results in a significant increase in astrocyte ramifications in the penumbra region as compared to the control cells (Fig. S2 I). These results are in agreement with those obtained from SMorph (Fig. 2 F). In conclusion, we found that SMorph could be used for accurate measurements of cell morphometry in a completely automated manner, and it was 1080 times faster than the ImageJ analysis, even when the faster of the two Sholl analysis methods was taken into account.

### Chronic antidepressant treatment produces subtle increase in parameters related to cell size

Studies investigating the glial density in depressed patients with varying histories of antidepressant usage relies on densitometric analysis of GFAP immunopositivity ^23,25^. Studies have consistently shown a decrease in GFAP immunopositivity in depressed patients ^23-25, 41^. Interestingly, antidepressant usage among depressed patients correlated strongly with the glial density ^23^. However, what contributes to the overall change in GFAP immunopositivity in 2-D images of brain sections is not clear. In order to assess whether chronic antidepressant treatment in mice induces structural remodeling in hippocampal astrocytes, we administered 20 mg/kg of desipramine i.p. for 21 consecutive days, while the control mice received equivalent volume (10 ml/kg) of 0.9% saline solution (Fig. 3 A). We first assessed various parameters related to cell size in 2-D projected images of fluorescently labeled GFAP-immunopositive astrocytes in the molecular layer of the upper blade of the hippocampal DG, where the afferents of perforant pathway form synapses (Fig. 3 B). We analyzed 50 cells in each group sampled evenly from 5 mice per group (Fig. 3 C). This region represents a major input area for the hippocampal formation. We found that chronic DMI treatment results in a significant increase in the 2-D projected surface area of astrocytes (Fig. 3 D). Furthermore, we also found a significant increase in the area of convex hull drawn encompassing the extremities of the GFAP-immunopositive portions of astrocytes (Fig. 3 E), suggesting an increase in cell territory. We further calculated the maximum radius of the cell as the distance from the cell soma to the farthermost end of a GFAP-positive branch. We found a significant increase in the maximum radius following chronic DMI treatment (Fig. 3 F). We further calculated the total length of the astrocytic branch skeletons and found a significant increase in the skeleton length upon chronic DMI treatment (Fig. 3 G). Interestingly, we also found a trend towards a decline in the average branch width of hippocampal astrocytes in DMI-treated mice (Fig. 3 H). Previously, it has been shown that chronic mild unpredictable stress (CMUS) results in a significant increase in the branch width, albeit in the hilar astrocytes ^27^. These results show that chronic DMI treatment and CMUS may have opposing effects on the thickness of astrocytic processes. Taken together, these results suggest that chronic antidepressant treatment results in an increase in cell size as inferred from 2-D images.

### Chronic desipramine treatment enhances branching in hippocampal astrocytes

We next assessed whether chronic desipramine treatment changes the properties of astrocyte branches. We found that the primary branches, that emanate directly from the cell body, significantly increase in number (Fig. 4 A). On the other hand, the average length of primary branches did not change upon chronic desipramine administration (Fig. 4 B). Similarly, we also found a significant increase in the number of secondary branches (Fig. 4 C) and a trend towards an increase in the number of tertiary branches (Fig. 4 E). However, the average length of secondary (Fig. 4 D) as well as tertiary (Fig. 4 F) branches did not alter significantly after the chronic desipramine treatment. We next calculated the number of branch points, and found a significant increase in the number of branch points in mice treated chronically with desipramine compared to controls (Fig. 4 G). We also found a significant increase in the number of terminal branches in desipramine-treated mice (Fig. 4 H). In summary, these data show that chronic desipramine treatment enhances the extent of branching in hippocampal astrocytes.

**Figure 4:**
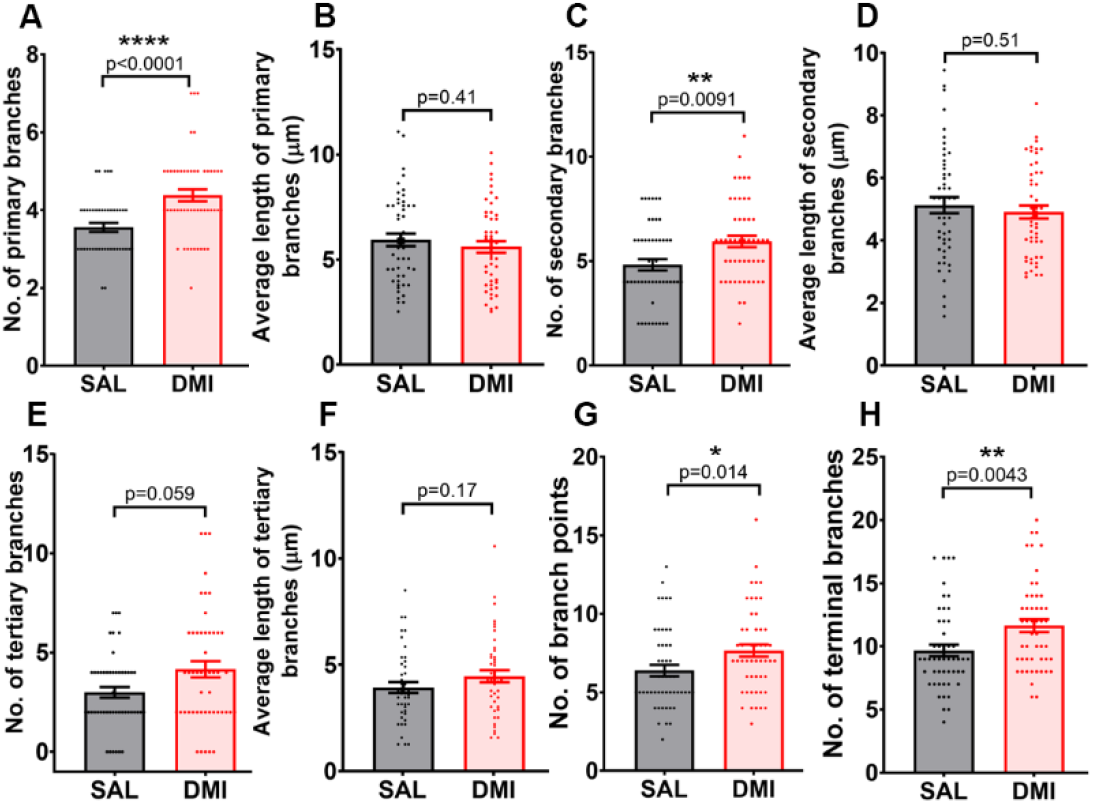
Chronic desipramine treatment enhances branching in hippocampal astrocytes. Parameters related to branching patterns were analyzed in saline and DMI-treated mice. Bar graphs with scatter plot shows a significant increase in the number of **(A)** primary, **(C)** secondary branches, as well as a trend towards an increase in the **(E)** number of tertiary branches in DMI-treated mice as compared to the controls. However, there is no difference between the two groups in the average lengths of **(B)** primary, **(D)** secondary or **(F)** tertiary branches. **(G)** Number of branch points and **(H)** number of terminal branches are also significantly increased in the DMI-treated group as compared to saline-treated controls. Bars represent mean±SEM. Asterisks represent statistically significant differences, with corresponding p values shown. n= 50 cells per group, evenly sampled from 5 mice per experimental group.

### Chronic desipramine treatment increases astrocyte ramification and results in structural remodeling of hippocampal astrocytes

We next performed Sholl analysis on the hippocampal astrocytes in saline and desipramine-treated animals. Sholl analysis showed a significant increase in the ramification of astrocytes (Fig. 5 A), with a significant increase in the area under the Sholl analysis curve (Fig. 5 B). We next calculated various parameters related to Sholl analysis and found a statistically significant increase in critical radius (Fig. S3 A), and a trend towards an increase in the critical value (Fig. S3 B). These correspond to the radius at which the local maxima of the polynomial regression occurs and the local maxima respectively. We also found a statistically significant decrease in the Sholl regression coefficient in the DMI-treated group (Fig. S3 C), signifying a slower rate of decay of the number of branches as a function of the distance from soma. There was also a statistically significant decrease in the Sholl regression intercept (Fig. S3 D). On the other hand, we did not see a statistically significant change in the ramification index, skewness and coefficient of determination (Fig. S3 E-G). Taken together, these results showed that chronic desipramine administration results in a significant increase in the ramification of hippocampal astrocytes.

**Figure 5:**
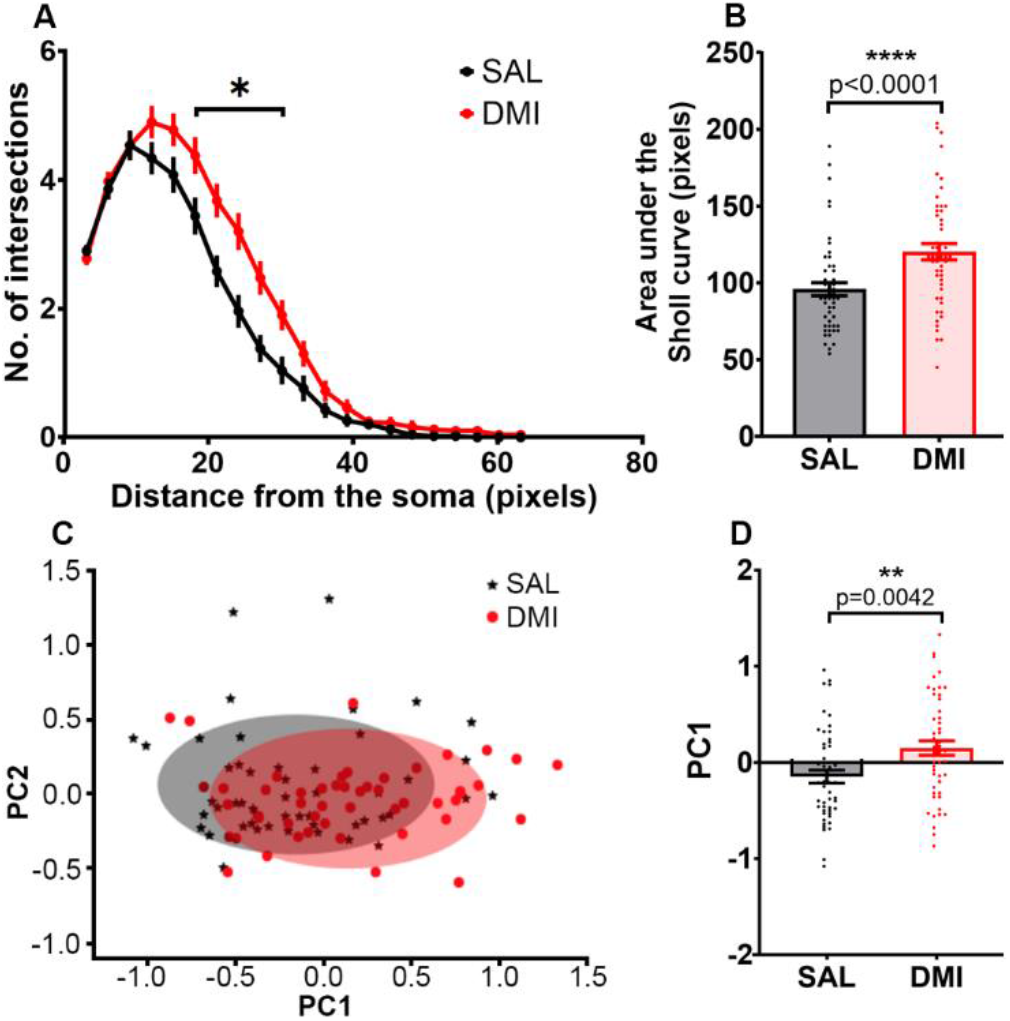
Chronic desipramine treatment increases astrocyte ramification and results in structural remodeling of hippocampal astrocytes. **(A)** Line plot of Sholl analysis reveals a significant increase in the astrocyte ramification in the DMI-treated group as compared to the controls. **(B)** Bar graph with scatter points shows a significant increase in the area under the Sholl analysis curve in the DMI-treated group as compared to the controls. **(C)** 2-D PCA plot representing the first two principal components, overlaid with ellipses corresponding to 3 standard deviations around the means are shown. **(D)** A bar graph with scatter points shows a significant difference in PC1 between the experimental groups. The line plot and bars represent mean±SEM. Asterisk in A represents discovery using the FDR approach. Asterisks in B and D represent statistically significant differences, with corresponding p values shown. n= 50 cells per group, evenly sampled from 5 mice per experimental group.

We next used PCA to reduce the dimensionality of the feature space comprising 23 morphometric features extracted from the images and Sholl analysis vectors of astrocyte images from saline- and desipramine-treated mice. The first two principal components described about 50% of the observed variability in the data (Fig. 5 C). The first principal component correlated positively with the complexity of astrocyte morphology and a t-test revealed a significant increase in the DMI group as compared with the saline group (Fig. 5 D). Taken together, these data show that chronic treatment with antidepressant desipramine results in significant structural remodeling in the hippocampal astrocytes. This reveals a novel kind of structural plasticity induced by antidepressant drugs in a non-neuronal cell type.

## Discussion

Study of changes in cellular morphology is an integral part of neurobiology research. Stimuli that influence the development or the onset or progression of neurodegenerative disorders robustly alter the morphology of neuronal and glial cells ^18,20,42^. On the other hand, subtler changes in morphology can be induced by physiologically relevant stimuli ^3-5,15,22,27,28^. Quantitative studies of cellular morphology *in vivo* have been challenging owing to the complex shapes of the cells of the nervous system and the time and efforts involved in such analyses. Here, we developed a software tool, called SMorph, to quantitatively analyze the morphology of cells in a rapid and completely automated manner. SMorph performs comprehensive morphological analysis taking into account multiple parameters related to cell size, branching pattern and radial distribution, thus revealing subtle morphological changes, which may not be accessible to traditional methods of analysis. SMorph also performs automated Sholl analysis. A large number of cells can be analyzed within seconds allowing analysis of large datasets. Furthermore, jupyter notebook provides an intuitive user interface, thus requiring no prior knowledge of python to perform the analysis.

In the comparison studies, SMorph took 40 seconds to complete the analysis and it was estimated to be 1080 times faster than the analysis in ImageJ, when the faster of the two Sholl analysis methods was considered. SMorph also computed much more additional morphometric information within that time, followed by PCA.

As a proof of principle, we analyzed the morphology of GFAP-positive astrocytes in the penumbra region around the stab wound in the *stratum oriens* of the hippocampal CA1. Matching ROI from the contralateral side was used to crop the control images of astrocytes. As expected, we found a significant difference in parameters related to cell size and branching. Interestingly, our analysis also revealed that the increase in branching is not uniform, but is more pronounced in the distal branches as compared to the proximal branches of astrocytes.

Furthermore, we wanted to test whether SMorph can be employed to detect subtle structural changes, unlike the visibly obvious changes seen after astrocyte reactivation. Emerging evidence implicates astrocyte atrophy as a putative pathogenic event in MDD. Additionally, long-term antidepressant use is associated with attenuated decrease in GFAP-immunopositivity in the hippocampus of MDD patients ^23^. Hence, we asked if chronic antidepressant treatment in mice alters the morphology of hippocampal astrocytes.

MDD is a debilitating neuropsychiatric illness and the currently available therapies are highly inadequate ^43^. A complete understanding of the pathophysiology of depression as well as the mechanisms underlying antidepressant action would help devise better therapeutic strategies. Conventional antidepressants require a long-duration treatment, before the behavioral effects appear ^44^. These antidepressants have also been shown to induce plasticity in neurons and neural progenitors in the hippocampal DG ^45-47^. Studies have shown that such plastic changes may be necessary for the behavioral effects of antidepressant drugs ^48^. However, it is not clear whether chronic antidepressant treatments also induce structural plasticity in non-neuronal cells.

Astrocytes have been shown to be affected in MDD patients ^23-25,41^, suggesting that astrocyte dysfunction may underlie the emergence of depressive behaviors. In particular, atrophy of astrocytes in the hippocampus may be therapeutically targeted. The density of hippocampal hilar astrocytes was reduced in the absence of antidepressant treatment in MDD patients ^23^. This decrease in astrocyte density was absent in the patients that received antidepressant treatment ^23^. Additionally, the area fraction of GFAP was inversely correlated with the duration of depression in the suicide victims ^23^. This suggests that astrocyte atrophy may be a potential pathogenic event in MDD, which may be reversed by chronic antidepressant treatment.

Studies in animal models have also shown that chronic stress results in reduced GFAP-immunoreactivity as well as reduced morphological complexity ^27-30^. It has been hypothesized that such degenerative changes in astrocytes as a result of chronic stress may play a causative role in the behavioral symptoms of depression. This hypothesis is further supported by the evidence showing that selective ablation of prefrontal astrocytes is sufficient to induce depressive-like symptoms in rats ^31^. Hence, reversing the astrocyte atrophy can be a promising therapeutic avenue to be explored.

It was recently shown that chronic mild unpredictable stress (CMUS) alters the morphology of hippocampal astrocytes and that this effect is most pronounced in the molecular layer of the DG ^27^. This prompted us to ask whether chronic antidepressant treatment, which is known to reverse the effects of chronic stress also alters the morphology of astrocytes in the molecular layer of the DG. In contrast to the stab wound, we didn’t observe visibly obvious changes in the morphology of astrocytes in the DMI-treated mice when compared to the controls. However, upon comprehensive quantitative analysis with SMorph, we found that the chronic DMI administration does induce morphological plasticity in the astrocytes of the DG molecular layer of the hippocampus.

Using SMorph, we extracted 23 morphometric features from 50 cells per experimental group from 5 mice per group. Comparative analysis showed a significant increase in parameters related to cell size and branching complexity.

Sholl analysis has been used extensively to study the morphology of cells with processes ^32-34^. However, most of the available tools require painstaking manual tracing of the processes. Although some methods do allow Sholl analysis without manual neurite tracing ^33^, they still work on one cell at a time. Additionally, they haven’t been tested for adaptability to various cell types in the brain. SMorph, which we have tested on neurons, microglia and astrocytes, performed automated Sholl analysis on 100 cells within seconds. Sholl analysis revealed a significant increase in astrocyte ramification in DMI-treated mice as compared to saline-treated controls. In addition, we found significant differences in low-dimensional features obtained using PCA. In future, further clustering of the low-dimensional PCA data using clustering algorithms may help cluster morphologically distinct subpopulations of astrocytes, which may be differentially affected by stimuli such as chronic antidepressant treatments. Moreover, It is becoming increasingly clear that astrocytes are not a homogeneous group of cells as previously thought, but are diverse in their gene expression, physiology and morphology ^49,50^. It is postulated that these differences are a result of the functions of the brain region that they are associated with ^49^. However, the presence of distinct functionally-specialized subpopulations of astrocytes within a brain region can not be ruled out. Assuming that the astrocytic structure is designed to optimize their function, multivariate analysis of astrocyte morphometry may help sort these cells into functional subpopulations. This would necessitate the multiparametric morphometric analysis such as that performed by SMorph, combined with imaging, electrophysiology and single cell transcriptomics ^50,51^.

Additionally, astrocytes associated with neurological disorders are also not homogeneous. It was shown that the reactive astrocytes associated with neuroinflammation and ischemia are polarized into distinct A1 and A2 subtypes respectively ^52^. While A1 reactive astrocytes are thought to be neurotoxic, A2 reactive astrocytes seem to be neuroprotective ^52^. Also, single cell RNAseq analysis revealed the presence of Alzheimer’s disease (AD)-specific populations of astrocytes that begin to appear early in the mouse models of AD^53^. In the wildtype controls, these cells appear only in the old age ^53^. Astrocytes with similar gene expression profiles were also observed in aged human brains ^53^, showing the enormous translational potential of such studies. It would be interesting to perform detailed multivariate morphometric studies in disease models to identify whether morphometry reveals disease-specific signatures. SMorph also works well on images of neurons and microglia, and in principle could be used to analyse the morphology of any cell type in the body with stellate morphology.

In conclusion, we believe that SMorph would help accelerate the research related to morphometric changes in the cells of the nervous system and beyond. It would also help quantify subtle changes in morphology induced by physiologically-relevant stimuli, which may have been inaccessible to conventional techniques. Indeed, we found that chronic DMI treatment induces significant morphological changes in hippocampal astrocytes. This data reveals a novel form of structural plasticity in a non-neuronal cell type induced by chronic antidepressant therapy. It would be interesting to understand whether such morphological plasticity in astrocytes is essential for the behavioral effects of chronic antidepressant treatments.

## Author contributions

PS wrote the code for morphological analysis, and performed data analysis. GV performed animal experiments, imaging, data analysis and helped with illustrations and writing, SCRT performed the stab wound surgeries. NR provided resources and expertise for the stab wound experiments and helped with writing. SM conceived the study, wrote the manuscript and obtained funding. All authors proofread the manuscript and approved the final version.

## Acknowledgements

This work was supported by the INSPIRE faculty grant from DST, India to SM and Early Career Research Award from SERB to SM. GV was supported by CSIR-NET Junior and Senior Research fellowships. SCRT was supported by N-PDF Post-doctoral fellowship from SERB, DST, India. NR was supported by the DBT-IISc partnership program and SwarnaJayanti Fellowship, DST, India. Authors would also like to thank Mr. Manjunath and the staff at the central animal facility and the bioimaging facility at IISc for technical help.

## Data Accessibility Statement

The source code is available in the supplemental zip file (SMorph.zip). Periodic updates to this code will be available at https://github.com/parulsethi/Morphometric-analysis. Requests for raw data can be addressed to the corresponding author and the information will be made available upon reasonable request.

---

**Figure S1:**
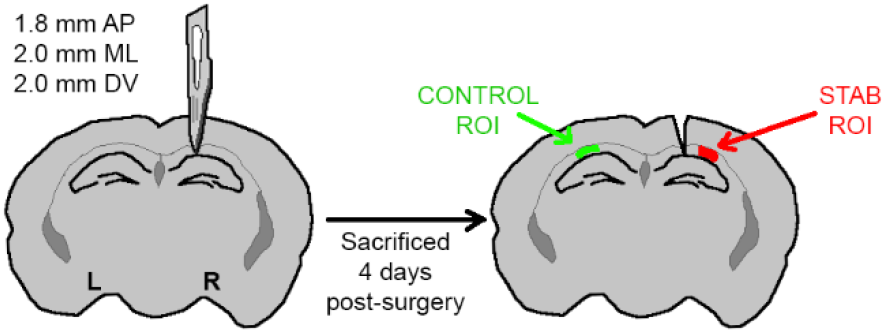
Schematic showing the stab wound location and ROI selection. Mice underwent a stab wound surgery with #11 scalpel blade targeting the hippocampal CA1 *stratum oriens* region. 4 days following the surgical procedure, mice were perfused by transcardial perfusion and the brain sections were immunostained for GFAP. a ROI in the penumbra regions slightly away from the wound was selected for cropping the astrocytes for the stab wound group (here shown in red). A matching ROI from the contralateral side was used as a control (shown in green). 3 mice were used for stab wound surgery.

**Figure S2:**
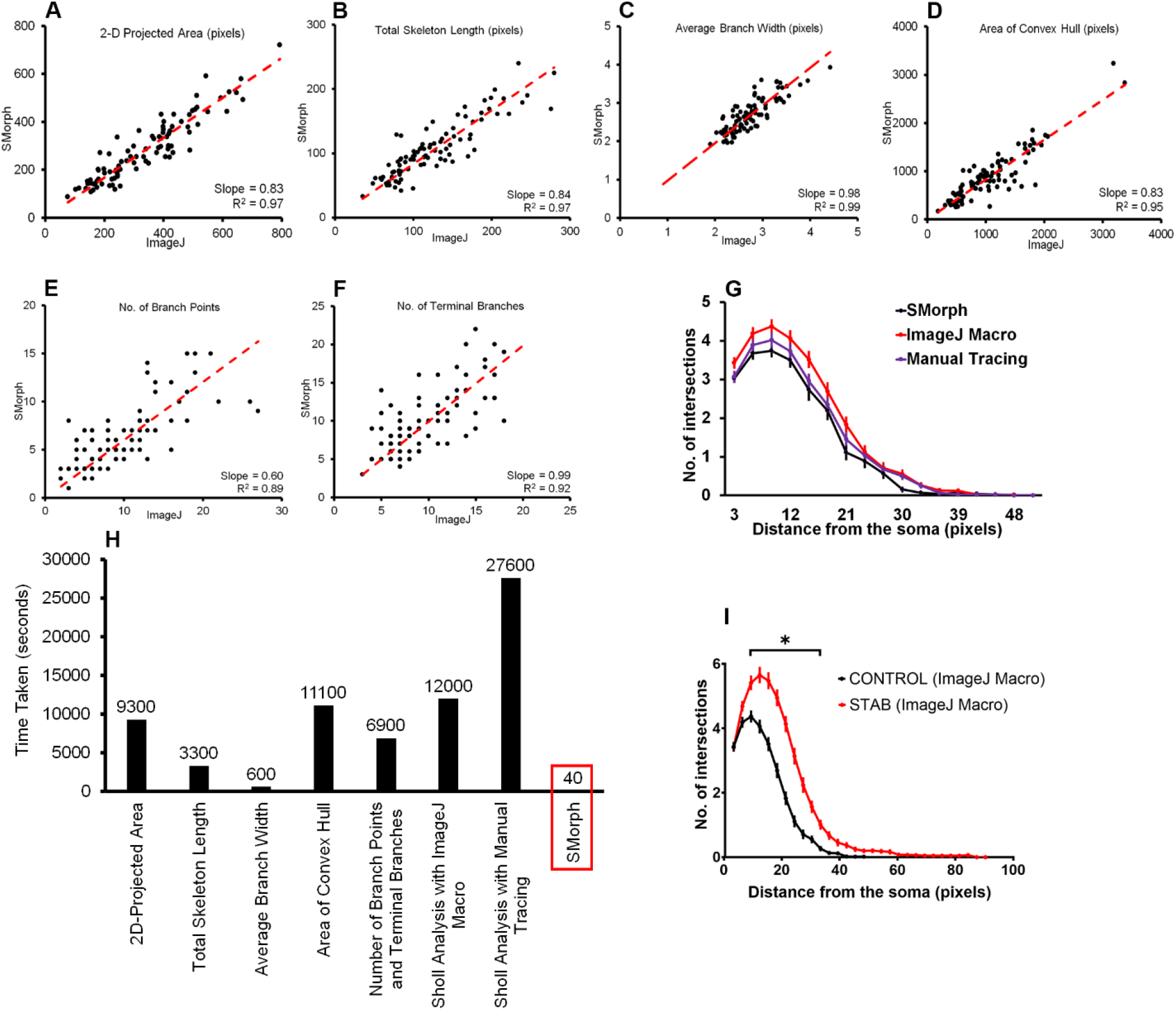
SMorph allows quick and accurate morphometry analysis. The performance of SMorph was compared against that of available tools in ImageJ. The linear regression through origin was performed on the 2 dimensional datasets from the two methods. Scatter plots show the corresponding values for individual cells, whereas the dashed red line represents the linear regression trendline for **(A)** 2D-projected area, **(B)** total skeleton length, **(C)** average branch width, **(D)** area of the convex hull, **(E)** number of branch points and **(F)** the number of terminal branches. The values of R^2^ and slope are also denoted. **(G)** The line plot shows the results of comparative Sholl analysis on the same set of cells with 3 different methods, namely, SMorph (black line), using unsupervised skeletonization using ImageJ Macro (red line) and using completely manual tracing (purple line). **(H)** Histogram depicts the time taken for various steps of analysis in seconds and is compared against SMorph. **(I)** The line plot shows Sholl analysis comparison between astrocytes in the penumbra region near the stab wound compared with the control cells on the contralateral side. Results in G and I are represented as mean±SEM. n=46 cells from 3 mice for A-H. n= 46 cells (controls) and 48 cells (stab) from 3 mice in I. Asterisk in I represents discovery using the FDR approach.

**Figure S3:**
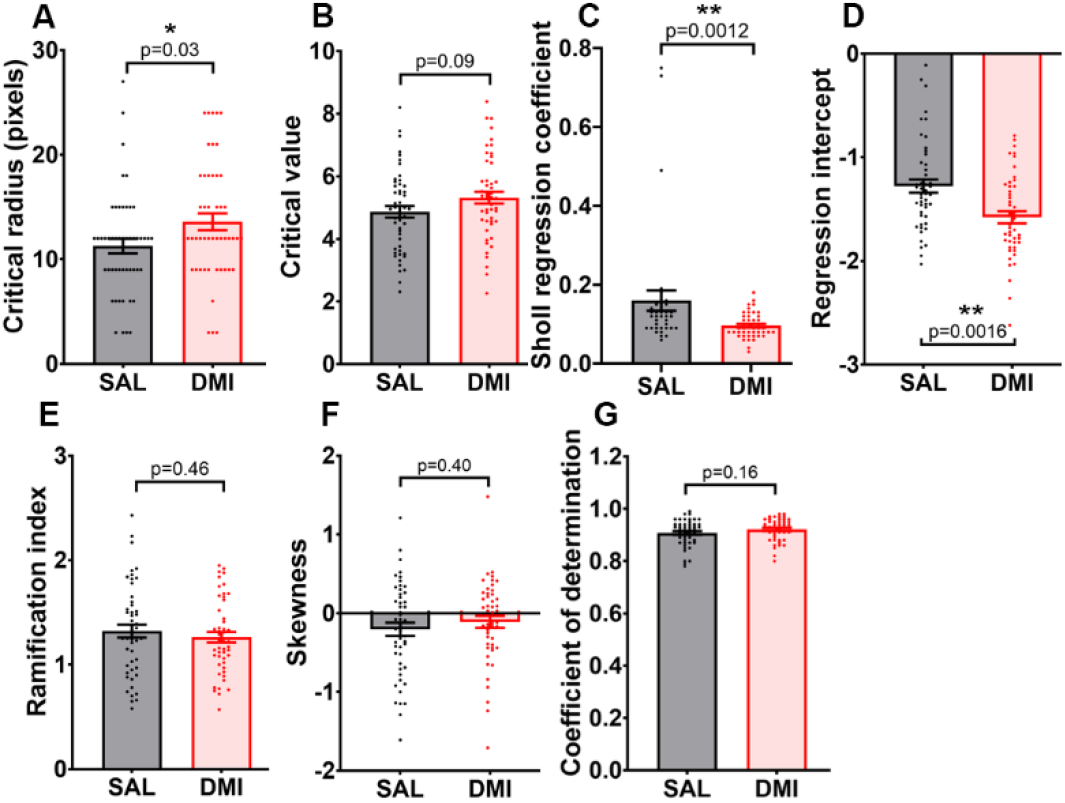
Parameters related to Sholl analysis show a significant change in astrocyte branching. Bar graphs with scatter points show **(A)** a statistically significant increase in the critical radius, **(B)** a trend towards an increase in the critical value and **(C)** a significant increase in the Sholl regression coefficient in DMI-treated astrocytes as compared to controls. **(D)** The bar graph with scatter points shows a statistically significant decrease in the Sholl regression intercept. However, there is no change in the **(E)** ramification index, **(F)** skewness and **(G)** coefficient of determination between the 2 groups. Bars represent mean±SEM. Asterisks represent statistically significant differences, with corresponding p values shown. n= 50 cells per group, evenly sampled from 5 mice per experimental group.

